# A vacuum-actuated microtissue stretcher for long-term exposure to oscillatory strain within a 3D matrix

**DOI:** 10.1101/149336

**Authors:** Matthew Walker, Michel Godin, Andrew E. Pelling

**Author notes:** Author for correspondence Andrew E. Pelling, 598 King Edward, University of Ottawa, Ottawa, ON K1N 6N5, Canada, Web: http://www.pellinglab.net.

## Abstract

Although our understanding of cellular behavior in response to extracellular biological and mechanical stimuli has greatly advanced using conventional 2D cell culture methods, these techniques lack physiological relevance. We developed the microtissue vacuum-actuated stretcher (MVAS) to probe cellular behavior within a 3D multicellular environment composed of innate matrix protein, and in response to continuous uniaxial stretch. The MVAS consists of an array of fifty self-assembled microtissues bordered by vacuum chambers. When a vacuum is applied, the microtissues stretch in plane allowing live imaging. The MVAS is highly suitable for biomedical research and pharmaceutical discovery due to a high-throughput array format and scalable fabrication steps outlined in this paper. We validated our approach by characterizing the bulk microtissue strain, the microtissue strain field and single cell strain, and by assessing F-actin expression in response to chronic cyclic strain of 10%. The MVAS was shown to be capable of delivering reproducible dynamic bulk strain amplitudes up to 13% and the strain field had local maxima around each of the cantilevers. The strain at the single cell level was found to be 10.4% less than the microtissue axial strain due to cellular rotation. Chronic cyclic strain produced a 35% increase in F-actin expression consistent with previously observed cytoskeletal reinforcement in 2D cell culture. The MVAS may further our understanding of the reciprocity shared between cells and their environment, which is critical to meaningful biomedical research and successful therapeutic approaches.

## Introduction

Cellular behavior is highly influenced by biological and mechanical stimuli from the surrounding extracellular environment.^1–5^ Whereas in the body cells receive stimuli in three dimensions, conventional cell culture techniques limit these interactions to two dimensions. In doing so, adhesion complexes and the cytoskeleton are forced into an unnatural apical-basal polarity.^6^ Conventional cell culture in petri dishes further disrupts adhesion-signaling pathways by offering an extremely rigid substrate void of natural matrix adhesion ligands, which greatly impacts cellular morphology and phenotype.^2,4^

In addition to a soft 3D matrix, many cells in the body also experience cyclic mechanical stretch. Examples include the expanding of airways during inspiration or of blood vessels during systole. These forces create continually unstable and unevenly distributed strain at focal adhesion complexes, across the cell membrane, along cytoskeleton filaments and through the nucleus. Many ligand-receptor affinities that regulate cellular phenotype and function are known to be responsive to these forces. ^1,7^.

The differences between the *in vivo* extracellular environment and a 2D static, rigid petri dish may account for observed disparities in cellular behavior and could explain how many drugs developed using conventional 2D cell culture lose their efficacy in costly clinical trials.^3,5,8,9^ Thus there is a need for novel high-throughput, low resource intensive cell culture techniques capable of probing cellular behavior and drug screening while providing a physiologically relevant environment.

To recapitulate the 3D *in vivo* environment, cells have often been cultured in bulk soft gels of innate extracellular matrix proteins. Due to their scale, these methods are resource intensive and low throughput, while leading to a high diffusive barrier for pharmacological treatments and nutrients. The ability to image through the sample is also often limited. The disadvantages of large-scale 3D cell cultures led to the development of sub-millimeter 3D cell culture models called microfabricated tissue gauges (microtissues)^10^. The microtissue model consists of an array of wells each containing two flexible polydimethylsiloxane (PDMS) cantilevers spaced approximately 500 μm apart. Cells in a collagen solution are introduced into each well, and self-assemble around the tops of the cantilevers into an organized 3D structure highly comparable to *ex vivo* tissue.

Recently, microengineered devices have been developed to deliver cyclic stretch to microtissues. For example in the Magnetic Microtissue Tester (MMT), a magnetic microsphere is manually fixed under a microscope with tweezers to one of the cantilevers in each well, and is pulled by magnetic tweezers.^11,12^ The MMT was later adapted to stretch multiple microtissues simultaneously with an array of electrodeposited bar magnetics.^13^ This method, however, still requires manually fixing the microspheres to one cantilever at a time, which limits high throughput fabrication, and the bulk strain amplitude is limited to 4%.

Here we present an alternative method, the Microtissue Vacuum-Actuated Stretcher (MVAS), which utilizes vacuum actuation to produce bulk strain amplitudes up to 13%, and is fabricated by scalable mold replication and alignment steps. It consists of an array of wells each with a set of cantilevers on a thin membrane bordered by vacuum chambers. When a vacuum is applied, the walls of the wells deform, stretching the thin membrane and the microtissue in plane allowing simultaneous live-cell imaging. Similar vacuum actuation systems have previous been shown to be reliable for 2D cell culture.^14–16^ In this paper we describe the fabrication steps of the MVAS, and to validate our approach, we characterize the bulk microtissue strain, the microtissue strain field and single cell strain, and assess changes in F-actin expression in response to chronic cyclic strain.

## Methods

### Device Design

The MVAS consists of five independently controllable rows of ten microtissue wells. As with the original microtissue design,^10^ each well has an open top for cell loading and cantilevers spaced apart by 500μm around which the microtissues are formed. The wells are bordered by enclosed vacuum chambers. When a vacuum, controlled through an external electronic regulator (SMC, ITV0010) via Labview software, is applied, the flexible PDMS material deforms, separating the cantilevers to stretch the microtissues along their longitudinal axes.

The MVAS is composed of a top support and three photolithographic layers all entirely polydimethylsiloxane (PDMS). An exploded view is shown in figure 1a) and the assembled device is shown in figure1b). The top support contains the vacuum and fluidics inlets, and encloses the fifty-microtissue wells within a single shared media well. During fabrication, the top support reduces shrinking and warping of the top layer so it retains the geometry of its mold. The top layer contains cell and vacuum chambers. The middle membrane is a thin layer with the cantilevers around which the microtissues compact. The bottom layer contains vacuum chambers with identical geometry to the top layer, and empty bottom chambers that equalize the pressure on either side of the membrane to minimize out of plane motion.

**Figure 1:**
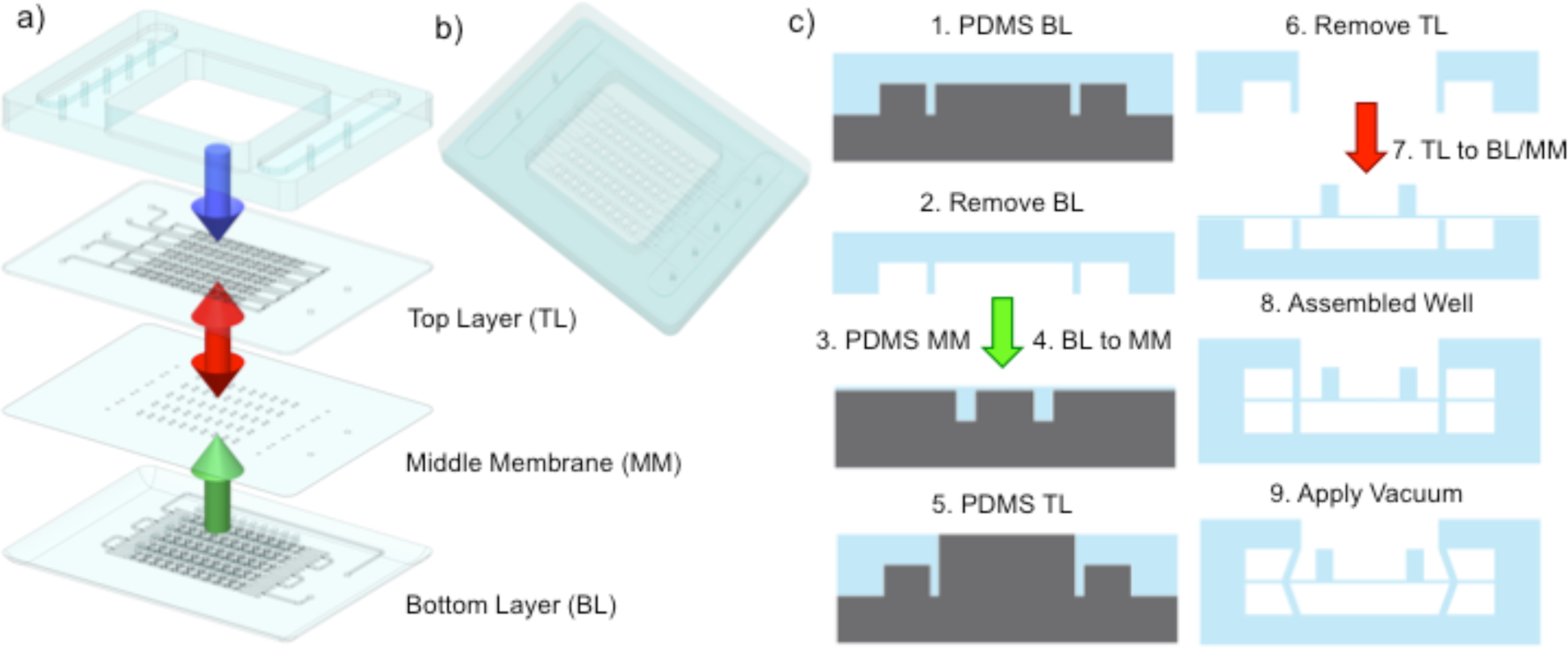
MTMS device assembly. The MVAS consists of 4 layers: 1) a top support; 2) a top channel and vacuum layer; 3) a middle membrane with the cantilevers; and 4) a bottom channel and vacuum layer. An exploded view is shown in a) and the assembled device is shown in b). A cross-section of fabrication steps of one well are illustrated in c). For the bottom layer, PDMS is applied over top of the features on the SU-8 photolithography master and removed. For the middle membrane and top channel layers, PDMS is spin coated to 30μm and up to the top of the features, respectively. The bottom layer is then plasma bonded onto the middle membrane (green arrow) and the top support is bonded onto the top layer (blue arrow). The two are then removed from the middle membrane and top layer masters, respectively, and bonded together (red arrow). When vacuum is applied, the middle membrane is strained in plane moving the cantilevers apart stretching the microtissue.

### Device Fabrication

The top layer, middle membrane, and bottom layer were fabricated by mold replication from photolithographic masters. Briefly, SU-2075 photoresist (Microchem) was spin coated onto plasma-cleaned silicon wafers (Universitywafers.com). The photoresist was then UV polymerized through a photomask (CAD Art Services Inc.), transferring the features of the photomask to the wafer. Non-polymerized photoresist was then removed with SU-8 developer (Microchem). Spin rates, baking, and energy exposure were kept consistent with manufacturer’s recommendations. Two photolithography steps were used for the top layer to create open top wells, and for the through-holes in the middle membrane. The top and bottom photolithographic layers were cast from the SU-8 master with a 10:1 monomer to curing agent ratio, whereas the middle membrane was cast with a 15:1 ratio.

Fabrication steps of one well are illustrated in figure 1c). PDMS was applied over the features on the bottom layer master. For the middle membrane and top layer, PDMS was spin coated to 30μm and up to the feature height, respectively. The bottom layer was removed from its master and plasma bonded onto the middle membrane (green arrow). The top support was bonded onto the top layer (blue arrow). The two halves of the device were then removed from the middle membrane and top layer masters, respectively, and bonded together (red arrows).

### Cell Culture

NIH3T3 (ATCC) and NIH3T3-GFP (Cedarlane, AKR-214) cells were cultured in DMEM with 10% fetal bovine serum (FBS), 50mg/ml streptomycin and 50U/ml penicillin antibiotics (all from Hyclone Laboratories Inc.), and maintained at 37°C with 5% CO_2_ on 100mm tissue culture dishes (Fisher) until 80-90% confluent.

### Tissue Fabrication

Microtissue fabrication was performed as described previously,^10,17^ with modifications. Briefly, the device was sterilized by three washes with 70% ethanol, and treated with 0.2% Pluronic F-127 (P6866, Invitrogen) for two minutes to reduce cell adhesion to the PDMS. 300,000 cells were resuspended in 1.5mg/ml rat tail collagen type I (354249, Corning) substituted with 1x DMEM (SH30003.02, Hyclone), 44 mM NaHCO_3_, 15 mM d-ribose (R9629, Sigma Aldrich), 1 % FB S and 1 M NaOH to achieve a final pH of 7.0-7.4. The cell-collagen solution was pipetted into the MVAS and centrifuged to load ~800 cells into each well. Once the excess collagen was removed, the device was transferred into the incubator for 15min to initiate collagen polymerization. Cell culture media was added and changed every 24 hours. To measure the cell strain within the microtissue, one NIH3T3-GFP cell was included for every 200 non-labelled 3T3 cells. For all other experiments, solely non-labelled cells were used.

### Imaging

To estimate the tissue strain field, phase contrast videos were captured on a TiE microscope (Nikon) at 10 frames per second and 8-bit resolution. All other images were acquired on a TiE A1-R laser scanning confocal microscope (LSCM) (Nikon). 12-bit images were acquired with standard LSCM configurations with appropriate laser lines and filter blocks. To assess morphology, microtissues were fixed *in situ* with paraformaldehyde for 10 minutes and permeabilized with Triton-X for 3 minutes. The actin cytoskeleton was stained with Alexa Fluor 546 Phalloidin (Fisher, A22283) and the nuclei were stained with DAPI (Fisher, D1306).

### Bulk Strain

The bulk axial microtissue strain was calculated from measurements of cantilever motion. The locations of the cantilevers were tracked using pattern matching in Labview. To be consistent with previous work,^12,13^ the bulk strain was defined as the percent change in the distance between the inner most edges of the cantilevers. This measurement, however, does overestimate strain values if the microtissues are not perfectly anchored at the inner edges of the cantilevers. The peak dynamic bulk strain was determined from the magnitude at the fundamental frequency of the Fourier transform.

### Tissue Strain Field

Local strains were estimated across microtissues while undergoing a 0.1Hz sinusoidal stretch. The inter-frame displacements were estimated at five-pixel spacing across a region of interest encompassing the microtissue using a four level-pyramid based Lucas and Kanada algorithm^18^ with sub-pixel precision and a window size of seventeen-pixels in Labview. The inter-frame strain tensor was calculated from the gradient of the displacement field after it had been smoothed with a LOWESS surface-fitting algorithm in Matlab. Starting at 0% strain, the inter-frame strain field was integrated to estimate the local total strain field.

### Cell Strain

To measure the cell strain within the microtissue, randomly distributed GFP labelled cells were imaged at 2 days post seeding immediately following static loading at vacuum pressures of 0, 30, 60, and 90kPa. Individual cell Feret lengths were measured in matlab using adaptive thresholding on maximum intensity projections of confocal stacks. Only cells between the cantilevers were used in the analysis.

The cell strain, ε, was defined as the change in the maximum Feret length, L, divided by the initial length irrespective of cell orientation (equation 1).

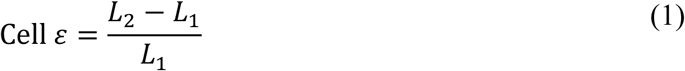

To account for reorientation of the cells caused by loading, the axial strain produced by cell lengthening and reorientation (Total Cell ε_x_) was calculated according to equation 2. The angle, θ, was measured between the cell Feret length and the longitudinal axis of the microtissue.

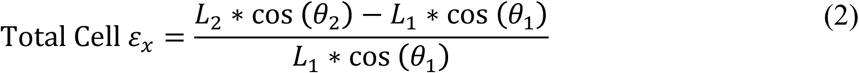

The contributions of cell lengthening (Cell ε_x_) and cell rotationn (Angle ε_x_) to axial strain were calculated according to equation 3 and 4, respectively.

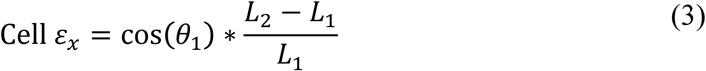

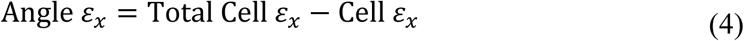

To compare the individual measurements of cell strain to the tissue strain, the axial tissue strain for each cell location was calculated from the gradient of the cell displacement field after it had been smoothed with LOWESS surface fitting algorithm.

### F-Actin Expression in Response to Chronic Strain

After microtissue fabrication, both the control and stretched groups were left to compact for two days under static condition. The stretched group then received a 10% 0.1Hz bulk strain for another two days while the control group was left under static condition. At four days post seeding, the nuclei and the F-actin cytoskeleton of both groups were stained with the same protocol, and imaged with identical laser and camera settings. The average integrated intensities of confocal stacks normalized to the control group were used as measures of cell number and actin polymerization. To assess the spatial distribution of changes in nuclei and F-actin, integrated confocal projections were aligned and averaged to give a percent change between stretched and control groups.

## Results

### Microtissue Morphology

Cells in a collagen solution were pipetted into the device, which was then centrifuged to load 770±30(SD, n=8) cells into each well. Microtissue formation occurred as been previously shown.^10^ The cells compacted the collagen matrix away from the F-127 coated sides and bottoms of the wells, and around the cantilevers into dense, organized freely suspended, three-dimensional microtissues. The MVAS contains fifty individual microtissue wells enabling high throughput manipulations into cellular mechanics and pharmaceutical discovery (figure 2a). At four days post seeding, microtissue survival rates were greater than 80%.

**Figure 2:**
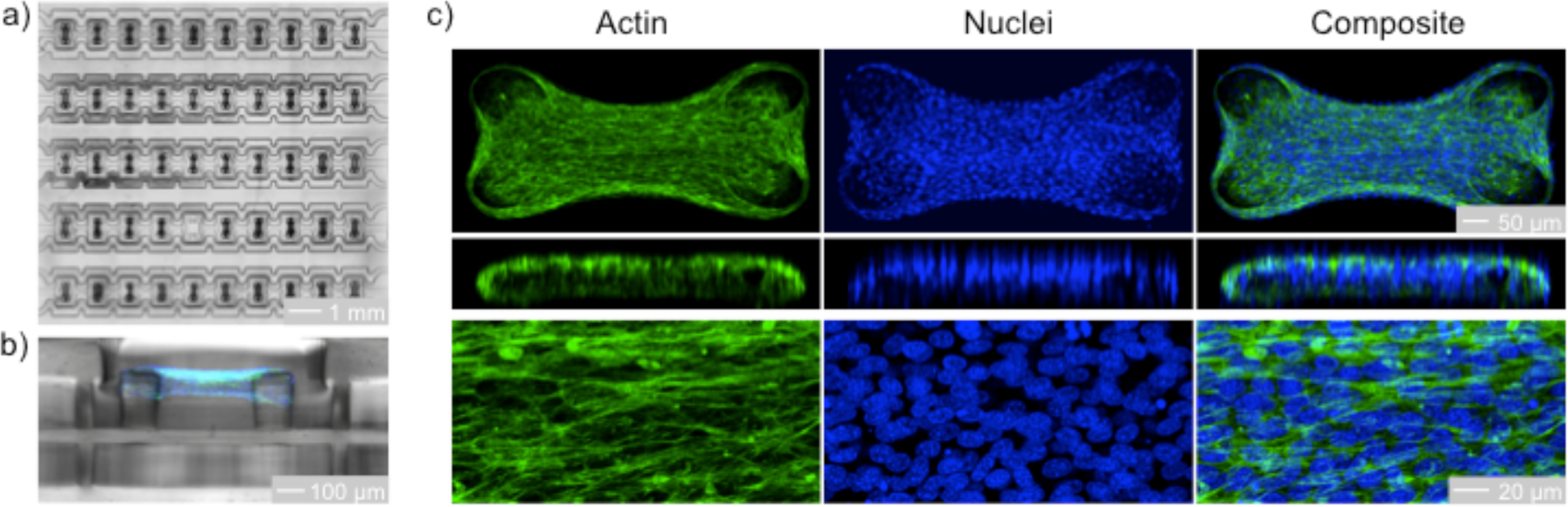
Representative microtissues at four days. Microtissues are dense, organized, three-dimensional cell cultures freely suspended around cantilevers. The MVAS contains an array of fifty microtissues (a). A cross-sectional view of the MVAS and a microtissue is shown in b). Max projections and an orthogonal slice of representative confocal stacks are shown in c). The actin cytoskeleton is in green and the nuclei are in blue. Both the actin cytoskeleton and nuclei possess a high degree of organization, aligning between the cantilevers.

A cross-sectional view of a fully compacted representative microtissue in the MVAS is shown in figure 2b). Maximum intensity projections of confocal stacks with an orthogonal slice at 10X magnification, and centrally-located maximum intensity projections of a 10μm-thick section at 60X magnification are shown in figure 2c). The actin cytoskeleton was highly polymerized and organized into dense stress fibers that were oriented with the length of the microtissue. The cell nuclei were evenly distributed in three dimensions and mostly aligned with the microtissue.

### Bulk Strain Characterization

When a vacuum is applied to the chambers bordering the cell culture wells, the cantilevers move apart, stretching the microtissues (figure 3a and movie 1). The motion is largely planar, allowing real-time imaging of the deformation. When using a computer-controlled regulator to apply a 0.1Hz sinusoidal vacuum, the bulk strain, measured from cantilever motion, was smooth and continuous with submicron resolution and high repeatability (figure 3b). The average (n=9) peak dynamic bulk strain was directly related to the applied vacuum with some nonlinearity at higher pressures (figure 3c). At 90kPa, the peak strain was 13.0 ± 0.9(SD)%, which covers the physiological range,^19–22^ and was highly reproducible.

**Figure 3:**
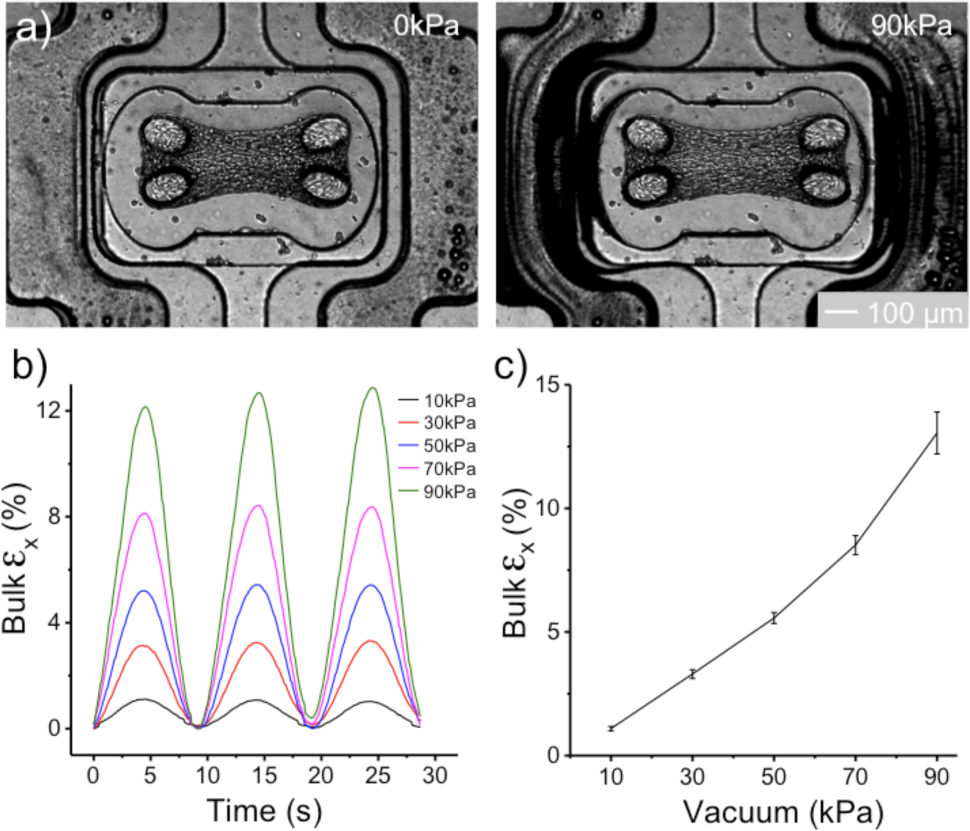
Bulk Strain Characterization. When a vacuum is applied, the cantilevers move apart stretching the microtissue in plane. Representative images of a microtissue in the MVAS are shown in (a). With a 0.1Hz sinusoidal vacuum, the bulk strain is smooth, continuous and repeatable (b). The average (n=9) bulk strain increased reproducibly with vacuum pressure (c). At 90kPa, the peak bulk strain was 13± 0.9(SD)%. Error bars represent the standard deviation.

### Tissue Strain Field

To estimate the dynamic axial strain distribution, microtissues (n=10) were stretched at 0.1Hz to a maximal bulk strain of 10.7±0.9(SD)%. As expected the bulk microtissue strain measured from the cantilever motion and the spatial average of the axial strain field followed the sinusoidally applied vacuum (figure 4a and b, respectively). The maximal spatially averaged strain field was and 8.3±0.7(SD)%. The difference in amplitude from the bulk strain can be accounted by an over estimation in the bulk strain measurement caused by imperfect anchoring of cells at the inner edges of the cantilevers.

**Figure 4:**
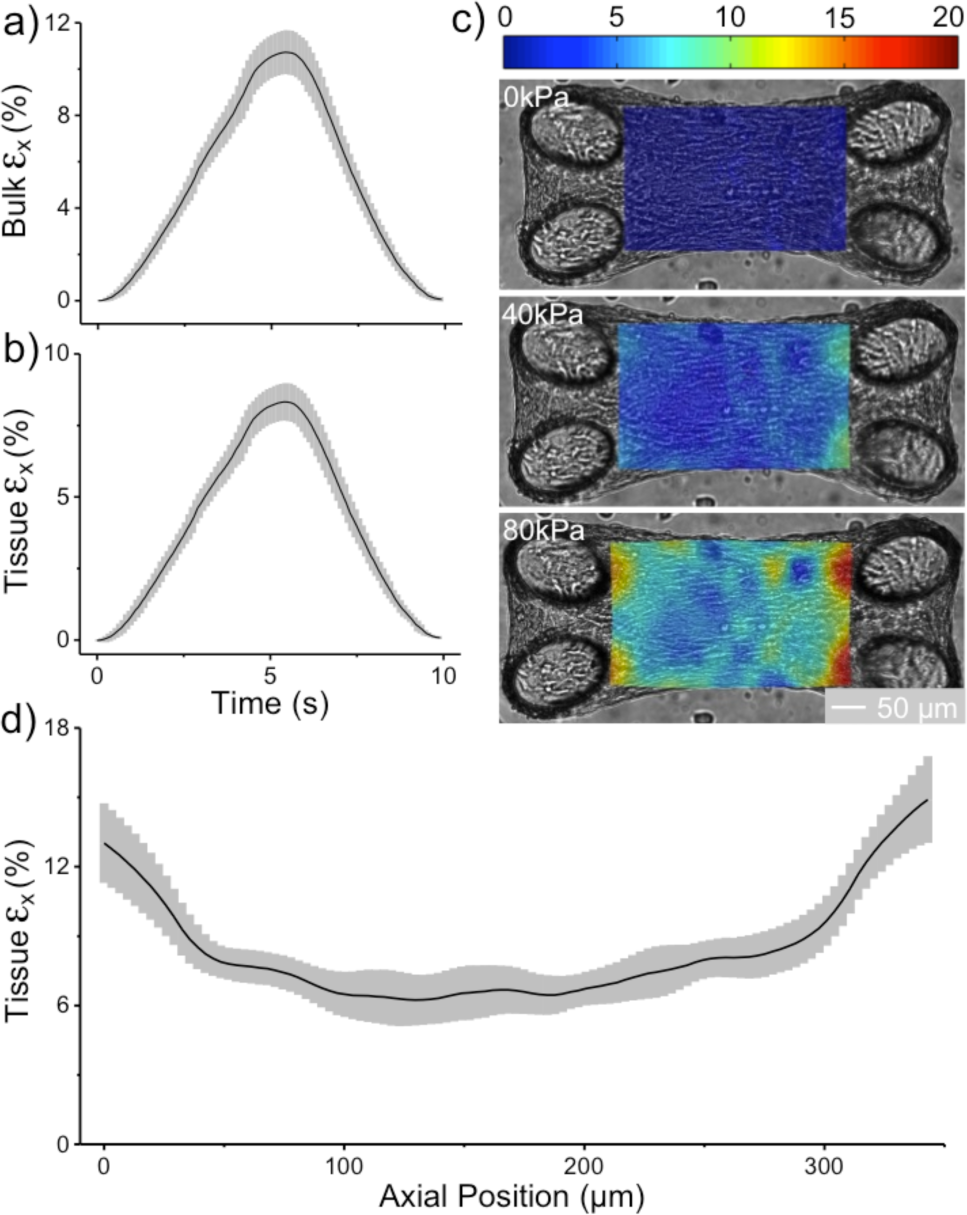
Tissue strain field. The time courses of tissue bulk strain measured from the inter-cantilever distance (a) and the average (n=10) of estimated axial strain fields (b) followed the sinusoidal change in vacuum. The estimated axial strain field of a representative microtissue is shown in c). The strain field was inhomogeneous and concentrated around the cantilevers. The average (n=10) peak axial strain values are plotted against axial position in d). The estimated strain field was highly reproducible between tissues. Error bars on all graphs represent the standard deviation.

Representative axial strain fields are shown in figure 4c) and movies 1-2. The transverse and shear strain fields were comparably smaller (movies 3-7). Qualitatively, there are local maxima in the axial strain field near each cantilever producing strain heterogeneity. The average (n=10) peak standard deviation of the axial strain field during one cycle was 2.8±0.4%.

The maximum axial strain during one cycle averaged across multiple (n=10) microtissues is plotted as a function of axial position in figure 4d). The strain heterogeneity produced by the local maxima near the cantilevers is again evident. The maximum axial strain field was reproducible across multiple microtissues with an average (n=10) standard deviation of 1.5±0.5%.

### Cell Strain

To measure the strain experienced by individual cells, changes in length of GFP labeled cells were assessed immediately following static loading within the MVAS (figure 5a). The average (n=79) cell length increased linearly (R^2^=0.999, p<0.001) with tissue axial strain measured from the gradient of the cellular displacement field (figure 5b). The degree of cell lengthening was significantly, albeit weakly, related to the initial cell length, with shorter cells undergoing greater strain (R^2^=0.07, p<0.05) (SI 1a)

**Figure 5:**
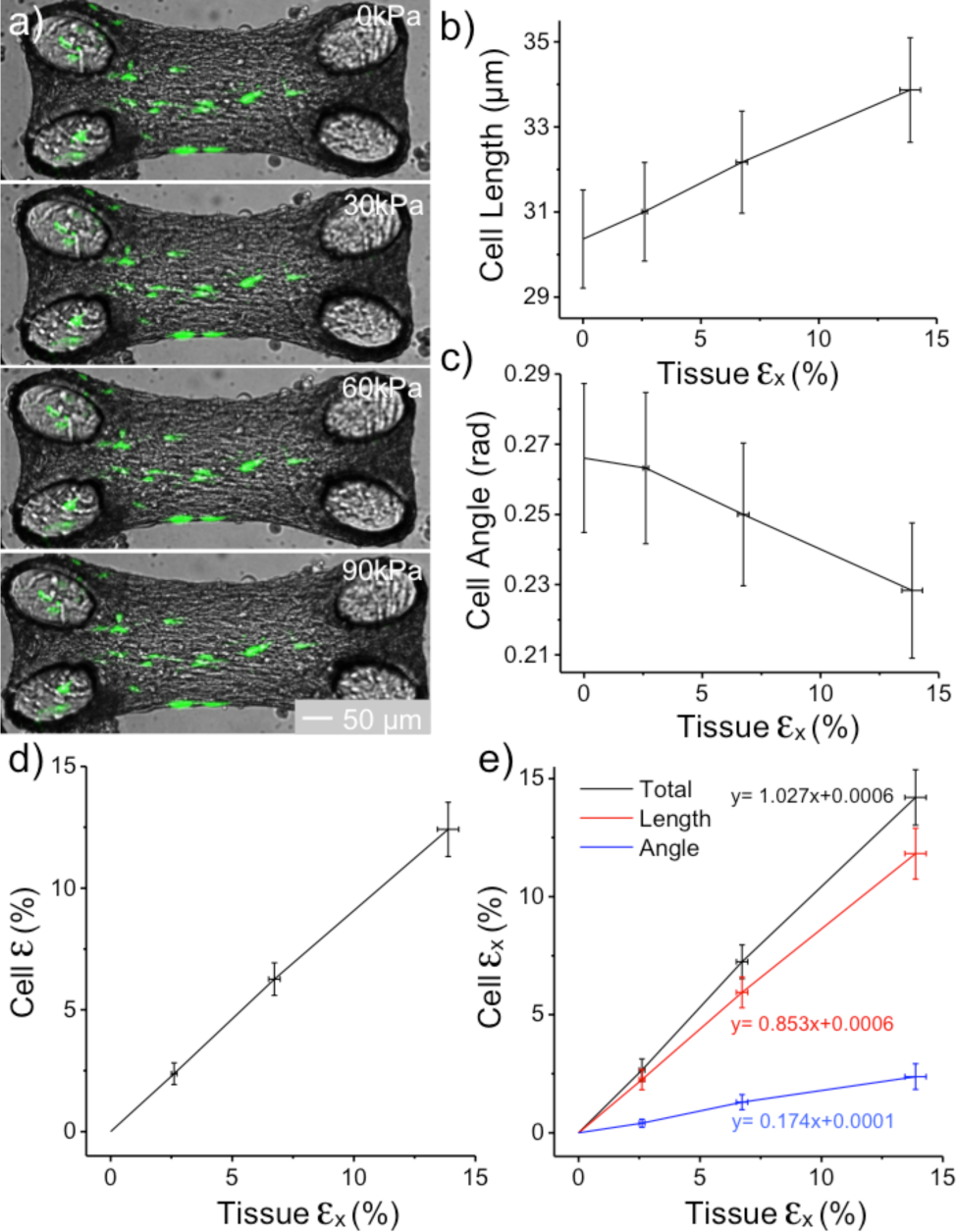
Cellular strain. A representative microtissue with GFP labeled cells (green) under various strains is shown in a). The average (n=79) cell length increased linearly with tissue axial strain (b) and the average absolute cell angle decreased (c). The average cell strain was 90% of the tissue axial strain (d) due to cellular reorientation. Accounting for reorientation, the axial cell strain was approximately equal to the axial tissue strain (e). The axial cell strain due to cell lengthening accounted for 85% of the axial tissue strain, and reorientation accounted for 17%. Error bars represent the standard error.

The average absolute angle between the cell and the longitudinal axis of the microtissue decreased (R^2^=0.987, p<0.005) with tissue axial strain (figure 5c). At a tissue strain of 13.9±0.4(SE)%, the average cell had reoriented 0.04±0.03(SE) rad to better align with the tissue. The degree of alignment was weakly related to the initial cell angle. Poorly aligned cells had a greater degree of reorientation (R^2^=0.26, p<0.001) (SI 1b). There were no relationships between initial cell angle and the cell or tissue strains (P>0.05) (SI 1c,d).

The average cell strain was 89.6% (R^2^=0.9996, p<0.001) of the tissue axial strain (figure 5d). This difference is attributable to cellular rotation (figure 5e). When accounting for rotation, the axial strain produced by cell lengthening and rotation was 102.7% (R^2^=0.9996, p<0.001) of the axial tissue strain. The axial cell strain due to cell lengthening accounted for 85.3% (R^2^=0.9996, p<0.001) of the axial tissue strain, and rotation accounted for 17.4% (R^2^=0.994, p<0.005). Interestingly, the relative contributions of cell lengthening and rotation to tissue axial strain were both constants at all strains tested.

### Chronic Stretch increases F-actin expression

To assess the effect of chronic stretch on F-actin expression, microtissues (n=6) were cyclically stretched at a bulk strain of 10.3±0.3(SE)% for 48 hours and compared to a static control group (n=6). The normalized integrated fluorescence of nuclei and F-actin is shown in figure 6a). Although there was no significant change in total nuclei fluorescence (p>0.05, t-test), chronic stretch significantly increased F-actin expression (p<0.01, t-test). On a per cell basis, F-actin expression increased 35±5(SE)%. The average nuclei and F-actin spatial distributions of percent change comparing stretched to control are shown in figure 6b) and c), respectively. Qualitatively, compared with the control, the nuclei appear slightly more concentrated in the center of stretched microtissues, and the F-actin fluorescence is greater throughout.

**Figure 6:**
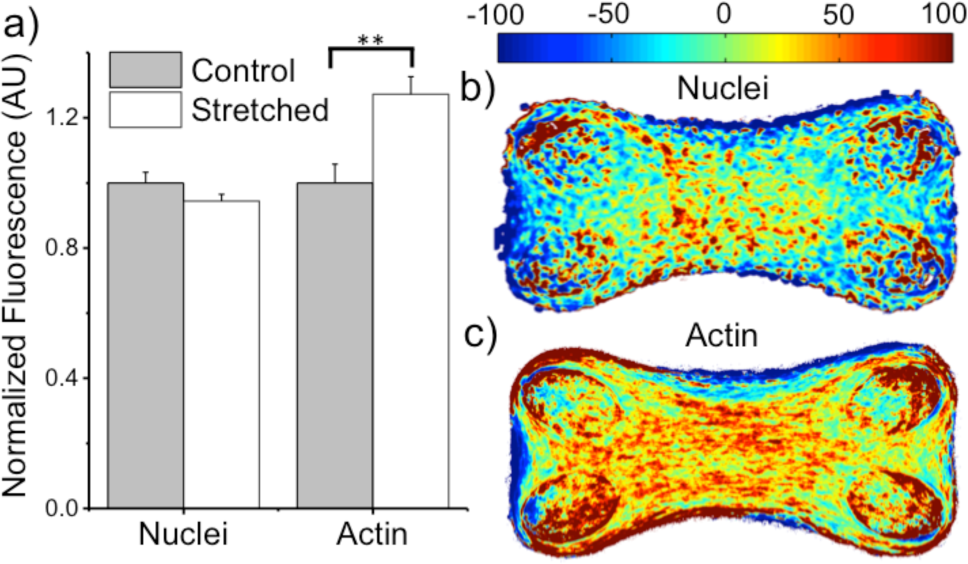
Cytoskeletal remodeling in response to chronic cyclic strain. The average (n=6) normalized total fluorescence of nuclei did not change between non-stretched control and stretched groups while F-actin fluorescence significantly increased with stretch (a). Chronic stretch caused a 35% increase in F-actin per cell. Error bars represent the standard error. The spatial distribution of percent change between stretched to control of average (n=6) nuclei and F-actin fluorescence are shown in b) and c), respectively. Stretch increased F-actin expression throughout the microtissues.

## Discussion

It is widely accepted that biological and mechanical stimuli from the surrounding extracellular environment influences cellular behavior.^1–5^ Yet traditionally, cell culture for biomedical research and pharmaceutical discovery has been carried out in petri dishes, which to a cell, is a 2D, rigid, static substrate with little physiological relevance. Although our understanding of cellular behavior has advanced tremendously using these conventional cell culture techniques, it is unknown whether cellular behavior studied in the lab truly reflects cellular behavior in the body. To answer this question, there is currently a need for cell culture methods with improved physiological relevance. To fulfill this need, cells in our device are cultured within a 3D collagen matrix and self-organize into a densely compacted microtissue comparable to *ex vivo* tissue. Importantly, cells grown in similar 3D collagen matrices better express their differentiated functions when compared to cells grown in 2D culture.^3,5,8,9^

In addition to offering relevant 3D cell-matrix interactions, our device enables investigations into chronic cyclic stretch. The uniaxial stretch is applied along the longitudinal axis of the microtissues, with which the cells are mostly aligned. The cellular orientation with respect to the strain direction recapitulates the circumferential stretch in airways or blood vessels, and the longitudinal stretch in muscle. We found chronic stretch significantly increases F-actin expression per cell. Cytoskeletal reinforcement in response to stretch is consistent with previously published findings in 2D culture. ^23–28^ Although the added complexity of a 3D matrix often makes direct comparisons with 2D culture difficult, it has been speculated that stretching 3D tissue constructs may induce more biomimetic effects than 2D cell culture.^29^

In addition to regulating F-actin, stretch has been shown in 2D culture to be a potent regulator of protein interactions within focal adhesion complexes that govern cell behavior.^19–22^ For two reasons it is critical to continue this work using 3D cell culture. First, a third dimension for cell adhesion significantly affects integrin/adhesion distribution and the cytoskeletal structure,^6^ which may alter how mechanical inputs are perceived. Second, the presence of a soft, viscoelastic matrix may alter the strain field felt by the cells. To that end, we found that the average cell strain was 10.4% less than the microtissue strain. However, after accounting for initial cell angle and rotation, the axial cell strain was equal to the axial microtissue strain. There are two possible explanations for this observation: 1) the matrix and the cells share the same modulus; or 2) microtissue loading is carried mainly through cell-to-cell junctions rather than through the matrix. Although future work is required to examine each of these explanations, we have demonstrated a viable method to stretch cells within a physiological 3D environment.

Biological and mechanical differences between 2D cell culture and the *in vivo* environment have cost pharmaceutical companies immensely in failed clinical trials creating a strong demand for more relevant 3D cell culture techniques.^3,5,8,9^ Although many large-scale bioreactors have combined a 3D cell matrix and stretch, these techniques often require months of cell culture, and due to their size, large drug doses to be synthesized.^8,30,31^ The multi-well capability and sub-millimeter scale makes the MVAS a high-throughput, low resource intensive alternative, and importantly, all fabrication steps are scalable to meet the demands of the pharmaceutical industry.

The MVAS is capable of delivering large oscillatory strain to an array of microtissues while allowing live imaging with minimal out of plane motion. Because of the planar stretch, the MVAS is especially suited for assessing cellular biomechanics. In this paper we quantified the bulk tissue strain, the tissue strain field and the cellular strain. The axial strain field was homogeneous apart from strain concentrations around the cantilevers. Strain gradients are important consideration for when examining local cell behavior^16^ and should be improved in future work through a better method of anchoring. Our device was capable of delivering a bulk tissue strain greater than 13% with a 90kPa vacuum. Although higher strain may be achievable through increased vacuum pressure or further optimizing device dimensions and materials, this strain sufficiently covers the physiological range in airways and blood vessels and is a sufficient stimulus to produce measurable differences in cell behavior.^19–22^

In this paper, we provided proof of concept for a device to stretch an array of lab grown microtissues consisting of fibroblasts and a reconstituted collagen matrix. Although this is a significant step forward in terms of physiological relevance compared to 2D cell culture, much is still needed to fully recapitulate the *in vivo* environment on a chip. Future work with using this device should be focused on culturing tissue specific cells, such as airway smooth muscle, vascular smooth muscle or skeletal muscle, and with a matrix composition that better matches health and disease states. To further the physiological relevance, each microtissue could be surrounded by a monolayer of epithelial or endothelial cells. With these future developments, combined with rapid gene expression and immunofluorescence techniques, the MVAS would be a leading platform for high-throughput chronic biomedical research and drug development in blood vessel, airway and muscle cell cultures.

## Summary

We developed a device to probe cellular behavior for biomedical research and pharmaceutical discovery while maintaining a physiologically relevant environment. Our device offers: 1) a soft, 3D multicellular environment composed from innate matrix protein; 2) the ability to apply large amplitude, long-term, cyclic mechanical stretch; 3) a micro-array format for high-throughput investigations; 4) compatibility with live imaging with limited out of plane motion; and 5) scalable fabrication steps. These features of our platform mark a significant improvement over other methods currently used to study cellular behavior in biomedical research and the pharmaceutical industry. Future investigations using the MVAS to chronically manipulate mechanical and biological cues may unveil how interactions between cells and their environment contribute to normal and pathophysiological behavior, which is critical to meaningful biomedical research and successful therapeutic approaches.

## Acknowledgements

M.W. is supported by OGS (Ontario Graduate Scholarship). The authors acknowledge support from individual NSERC Discovery Grants (M.G. and A.E.P.). A.E.P also acknowledges generous support from the Canada Research Chairs program.

